# Forensic likelihood ratios for shotgun sequencing data from DNA mixtures

**DOI:** 10.64898/2026.09.28.755155

**Authors:** Rasmus Nielsen, Abigail Daisy Ramsøe, Martin Sikora, Eske Willerslev, Thorfinn Sand Korneliussen

**Affiliations:** Department of Integrative Biology, University of California, Berkeley, Berkeley, CA, USA; Departments of Statistics, University of California, Berkeley, Berkeley, CA, USA; Center for GeoGenetics, University of Copenhagen, Copenhagen, Denmark

**Keywords:** Forensic Genetics, DNA mixtures, likelihood ratio test

## Abstract

Shotgun sequencing of DNA mixtures has the potential to become an important tool in forensic genetics, particularly for degraded samples or complex mixtures. However, the analysis of shotgun-sequenced mixtures is challenged by several confounding factors, including sequencing errors, population structure, the potential presence of relatives, linkage, and linkage disequilibrium. Additionally, the statistical methods currently available for such data are limited. Here, we develop likelihood ratios that summarize the evidence for the presence of a suspect’s DNA in a mixture, assuming genotypes of the suspect and the victim are available, while accounting for DNA from unknown contributors modeled either as general population contamination or as a single contaminating individual. We show that these likelihood ratios are not robust to model assumptions and can falsely support the presence of the suspect’s DNA when the true contributor is even a distant relative. We therefore propose a combined evidence score, Λ: the likelihood ratio minimized over contamination models, reported only when the evidence favors the suspect over a hypothesized sibling by at least a factor of C (default *C* = 10) under at least one contamination model, and otherwise set to 1 (log Λ = 0). In simulations based on 1000 Genomes data, Λ retains nearly all power to detect true contributors at mixture proportions as low as 5% while eliminating the false positives produced by model mis-specification, population structure, and by untested relatives. The methods are implemented in the open-source program DNAMIXTURE, providing the forensic and population genetics community a resource for robustly identifying small individual contributions to a DNA mixture.

## Introduction

The vast majority of forensic genetic analyses in the United States and beyond, are carried out using the standard 20 CODIS Single Tandem Repeat (STR) loci, or subsets thereof. There are good reasons for this. These markers are approximately unlinked (but see (Burri et al. 2015)), are highly informative, typically leading to high exclusion probabilities, and are thoroughly validated. However, Single Nucleotide Polymorphisms (SNPs) have emerged as attractive alternative markers in forensic genetics, particularly for degraded or low-quantity DNA samples (Mandape et al. 2024; Budowle et al. 2025). While SNPs are less informative per individual marker compared to STRs, their abundance across the genome enables simultaneous high-throughput analysis of millions of markers through SNP genotyping platforms or shotgun sequencing (Mandape et al. 2024; Ballard *et al*. 2020). An additional advantages of using SNP data is that these data can be generated using shotgun sequencing in situations where genotyping of STR markers fail due to DNA degradation (Pedroza Matute and Iyavoo 2025). Indeed, likelihood-based identification from extremely low-coverage shotgun sequencing data has been demonstrated (Vohr et al. 2015), and dedicated forensic SNP marker sets are now available (Tillmar et al. 2021). SNPs are, therefore, particularly valuable for recovering genetic information from evidence that would yield incomplete or no STR data (Budowle et al. 2025).

Despite the potential of SNP data generated using short-read sequencing, adoption remains limited due to high costs, technical complexity, and lack of standardized protocols and legal frameworks (Pedroza Matute and Iyavoo 2025). A distinct drawback is also the lack of appropriate statistical analysis methods (Cho *et al*. 2014). The analysis of SNPs is complicated by the error structure of the data, which needs to be modeled, and perhaps more importantly, the challenges that arise when using dense maps of SNPs in linkage and Linkage Disequilibirum (LD). Statistical evidence in court is typically presented as exclusion probabilities or as likelihood ratios, which compare the probability of DNA evidence under competing hypotheses (Balding and Nichols 1994; Evett and Weir 1998). In these calculations, it is generally assumed that markers are statistically independent of each other, i.e. that they are not in LD. However, both linkage between the markers and population structure will generate LD (Nei and Li 1973). Furthermore, linkage among markers can make it challenging to calculate accurate likelihood ratios distinguishing DNA contributions from relatives who will share long haplotypes, even if the chosen markers are not in LD in the general population.

Population structure is a challenge because of its role in generating LD, but it is also a substantial challenge for two other reasons. First, accurate likelihood ratio calculations require the availability of databases that are representative of the general population. While such forensic databases have been generated in many countries for the standard STR markers, they are typically lacking for SNP markers. Secondly, likelihood ratio calculations require the assumption of Hardy-Weinberg Equilibrium (HWE), or a correction for deviations from HWE must be included (Balding and Nichols 1994, 1995; National Research Council Committee on DNA Forensic Science: An Update 1996). Recent advances attempting to address some of these challenges for SNP makers include dynamic SNP selection methods and likelihood ratio formulas that account for sequencing errors in shotgun sequencing data (Andersen et al. 2025). SNP selection, as proposed by for example (Andersen et al. 2025), may at least partially address issues regarding LD. However, they do not address the issue of linkage, for example, if a suspect and the perpetrator are distantly related and share haplotypes that are longer than the typical extent of LD.

The analysis of DNA mixtures from crime scenes presents distinct challenges. For STR markers, the complexity of mixed profiles has necessitated increasingly sophisticated statistical methods (Haddrill 2021; Tao et al. 2018) with probabilistic genotyping software becoming widespread (Gill et al. 2021). The two most commonly used methods are, arguably, STRmix and TrueAllele, which employ Markov Chain Monte Carlo simulation to assign probabilities to possible genotype combinations (Thompson 2023; Bauer et al. 2020). These methods can produce dramatically different likelihood ratios for the same data(Thompson 2023). Recent comparative studies show that while both systems produce highly correlated results overall (greater than 88% correlation of likelihood ratios), differences become more pronounced for low-template contributors below 100 picograms (Greenspoon *et al*. 2024). More generally, there is substantial variation in mixture interpretation across laboratories (Butler et al. 2018), and a recent systematic review concluded that the scientific foundations of complex-mixture interpretation remain incompletely established (Butler et al. 2024). DNA mixture, therefore, remain a substantial challenge for STR markers. However, the increased resolution of short-read sequencing data may help resolve this issue. For analyses of such data, Crysup and Woerner (Crysup and Woerner 2022) introduced a read-level likelihood of two-person mixtures and used it to deconvolve mixtures into genotype posteriors for downstream imputation and kinship estimation (see also (Woerner et al. 2024, 2026) for mixture detection and mixture-fraction estimation from shotgun data). These methods recover genotypes but they do not provide likelihood ratios for forensic questions, and relatedness enters only through downstream analyses of the deconvolved genotypes rather than through competing hypotheses evaluated in the likelihood itself - which is our focus here. Related likelihood ratio methods have been developed for identification from low-coverage single-source samples. Nguyen *et al*. (2023) used likelihood ratios based on identity-by-descent to test whether sequencing reads originate from a known individual. Ouerghi *et al*. (2025) showed that this test in effect assumes, under the null hypothesis, that the person who truly deposited the DNA is included in the reference database used by the method, and that the resulting likelihood ratios can consequently overstate the evidence for identity by orders of magnitude.

The direct sequencing of individual haplotypes, and the availability of millions of markers, will ameliorate the statistical problems associated with the analysis of DNA mixtures of STR markers. However, the increased sensitivity of modern DNA methods creates a double-edged sword as many samples contain mixtures of minute DNA amounts from multiple contributors (National Institute of Standards and Technology 2019; Haddrill 2021). Analyses of such mixtures require statistical methods that account for this complexity.

In this paper we will describe and test statistical methods for analyzing mixtures of DNA in short-read sequencing data that will address some of these challenges. We will pay particular attention to the scenario where a sample has been obtained from a crime scene and genome sequencing has been carried out for both the suspect and the victim. The statistical challenge is then to determine if the sample contains DNA from the suspect, in addition to DNA from the victim and DNA from members of the general population.

## Methods

### Likelihood ratio of suspect contribution

In the following we will describe likelihood functions appropriate for short sequencing reads from mixture of DNA. We will assume the standard read nucleotide likelihood model:

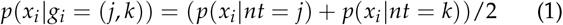

Here *x*_*i*_ is the observed read nucleotide in position *i* of the read. *nt* is a nucleotide (*nt* ∈ *S* = *A, C, T, G*) and *g*_*i*_ is a diploid genotype consisting of a pair of nucleotides, *g*_*i*_ ∈ *G* = *S*^2^. *p*(*x*_*i*_ | *nt* = *j*) is the usual rescaling of the phred-scaled read quality score (see. (McKenna *et al*. 2010; Li 2011; DePristo *et al*. 2011; Nielsen *et al*. 2011; Korneliussen *et al*. 2014)). For degraded DNA, damage can be incorporated into this calculation as in references (Briggs *et al*. 2007; Zhao *et al*. 2025).

We will assume a mixture of up to 3 different components: DNA from the victim, DNA from suspect, and DNA from the general population (contaminant). We will also, for now, assume linkage equilibrium among SNPs, HWE in the general population, and the availability of a reference panel providing accurate estimates of population allele frequencies. The full likelihood, for nucleotide *x*_*i*_ in position *i* of the read is then

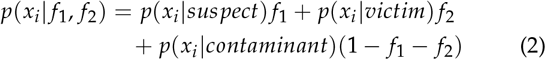

where *f*_1_ and *f*_2_ are the proportions of DNA from the suspect and the victim, respectively, *f*_1_ + *f*_2_ *≤* 1. Also, *p*(*x*_*i*_ | *suspect*) is the read nucleotide likelihood given by Equation (1) calculated for the suspect genotype and *p*(*x*_*i*_ | *victim*) is the similarly calculated read nucleotide likelihood calculated for the victim. Finally,

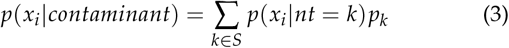

where *p*_*k*_ is the frequency of nucleotide *k* in the general population and *p*(*x*_*i*_ | *nt* = *k*), as before, is the standard rescaling of the read quality score. The total read likelihood is then obtained by taking the product of *p*(*x*_*i*_ | *f*_1_, *f*_2_) among all sites in the read and among all reads.

The expression above assumes that the contaminant reads are sampled independently of each other. That is approximately true if the number of people contributing to the contamination is vastly larger than the average number of reads per site. However, reads are not independent of each other when sampled from a limited number of individuals. The most extreme case is when contamination comes from a single person. In that case, the likelihood calculation becomes slightly more complicated because we have to sum over the possible genotypes of the contaminant individual:

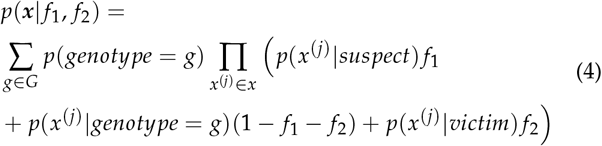

where ***x*** = {*x*^(1)^, *x*^(2)^, …, *x*^(*n*)^} is the set of the *n* observed reads in the position and, as previously, *p*(*genotype* = *g*) is calculated assuming Hardy-Weinberg equilibrium, e.g. *p*(*genotype* = (*AC*)) = 2*q*_*A*_*q*_*C*_ and 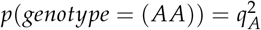, where *q*_*v*_ is the population allele frequency of nucleotide *v*. The index for the position has been suppressed in the notation. This expression can naturally be extended to multiple people, and perhaps it could be attempted to estimate the number of contaminating individuals. The expression for general contamination, and for a single individual contaminating, could be used at bounds on the LR under the assumption that these two extreme scenarios will bound the likelihood ratio under other scenarios of contamination. These equations, when taking the product over all SNPs, can be used to provide likelihood ratios by optimizing the likelihood for *f*_1_ and *f*_2_, and for *f*_2_ assuming *f*_1_ = 0. The resulting likelihood ratio presents the weight of the genetic evidence in favor of a contribution of the suspect versus no contribution of the suspect:

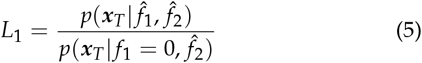

where ***x***_*T*_ represents that data for all reads in all sites (total data), with the subscript not to be confused with the previous index for position in a read.

There are substantial challenges associated with the presentation of this likelihood ratio directly in court. First, it is derived using strong assumptions regarding representative reference databases, HWE, and linkage equilibrium. We will address these challenges subsequently. However, we note a perhaps even more serious problem. With dense marker maps, even if the assumption of linkage equilibrium is approximately satisfied among markers, a relative could be part of the mixture causing long distance correlation among markers and spurious evidence of a contribution of the suspect. The difficulty in differentiating between a suspect and a close relative has been recognized since the early days of forensic DNA analysis (Evett 1992). The challenges in distinguishing relationship classes from genetic data, particularly in the presence of population structure, has been characterized in the context of familial searching (Bieber *et al*. 2006; Rohlfs et al. 2012). Dense SNP data exacerbate the problem: with genome-wide markers, even distant relatives of the true contributor can produce strong apparent evidence, as we show below.

We suggest addressing this problem by additionally using likelihood ratios that compare the evidence of a contribution of the suspect versus the evidence of a contribution of the suspect’s hypothesized sibling or parent. In other words, when arguing in court for a contribution of the suspect, we should evaluate how much evidence there is that, in fact, it is the suspect that contributed and not one of their close relatives. As we will show, presenting such likelihood ratios also generates some robustness to other violations of the assumptions of the model.

### Likelihood ratios of relatives

We can calculate the likelihood associated with a contribution from a hypothesized sibling as:

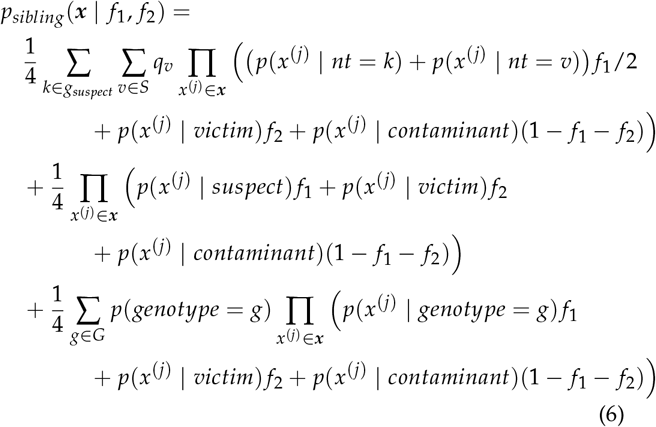

where, as previously, ***x*** = {*x*^(1)^, *x*^(2)^, …, *x*^(*n*)^} is the set of the *n* observed reads in the position, *q*_*v*_ is the population allele frequency of nucleotide *v, p*(*genotype* = *g*) is calculated assuming Hardy-Weinberg equilibrium, e.g. *p*(*genotype* = (*AC*)) = 2*q*_*A*_*q*_*C*_ and 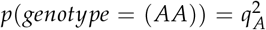. The index for the position has again been suppressed in the notation. This function then needs to be optimized similarly to the optimization for the function involving the suspect. The likelihood ratio is then

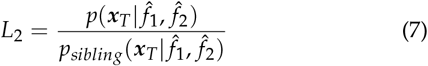

For a parent, the calculation is simpler:

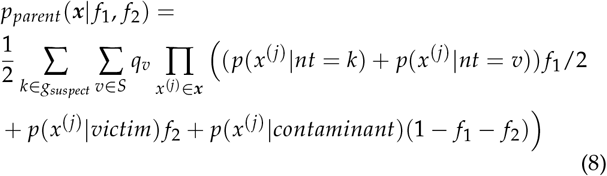

and the likelihood ratio is similarly

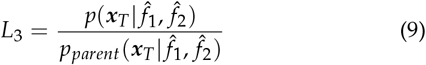

More distant relationships can be modeled in the same framework. Consider a *k*th cousin of the suspect (*k* = 1, 2, 3 for first, second, and third cousins). For such unilineal relationships the pair can share at most one allele identical by descent (IBD), and the probability of sharing exactly one allele IBD is *κ*_*k*_ = 2^*−*2*k*^, i.e. *κ*_1_ = 1/4, *κ*_2_ = 1/16, and *κ*_3_ = 1/64 for first, second, and third cousins, respectively. Conditional on the cousin sharing one allele IBD with the suspect, the likelihood is identical to the parent likelihood of Equation (8). Conditional on sharing no alleles IBD, the cousin is simply an (unknown) individual unrelated to the suspect, contributing a fraction *f*_1_ of the mixture:

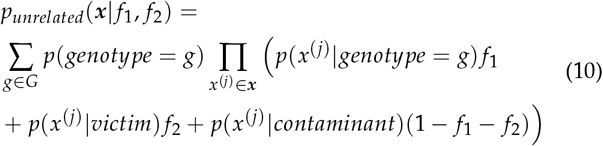

which is recognized as the last term of Equation (6). Averaging over the two IBD states, the likelihood for a hypothesized *k*th cousin is

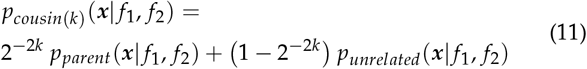

and the corresponding likelihood ratio is

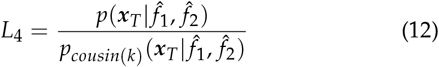

where, as for *L*_2_ and *L*_3_, the parameters are optimized separately under the numerator and denominator models. We note that Equation (11) applies more generally to any relationship in which the relative and the suspect can share at most one allele IBD, by replacing 2^*−*2*k*^ with the appropriate probability of sharing one allele IBD: 1/2 for half-siblings, avuncular pairs, and grandparent–grandchild pairs, and 2^*−*(2*k*+1)^ for *k*th cousins once removed. In this notation the sibling likelihood of Equation (6) is likewise the mixture 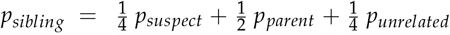, where *p*_*suspect*_ denotes the likelihood of Equation (2) with the suspect as contributor, reflecting the sibling IBD-state probabilities (1/4, 1/2, 1/4) for sharing (2, 1, 0) alleles. As in the case of *L*_1_, versions of these expressions assuming a single contaminating individual are obtained by additionally summing over the genotype of the contaminant as in Equation (4).

### Numerical considerations

The likelihood functions described above need to be evaluated numerically and optimized. It includes sums over products which can be evaluated numerically using standard methods employing log sums and scaling factors. The optimization is either one-dimensional or two-dimensional and can easily be achieved using standard algorithms. We distribute a C program, DNAMIXTURE, which uses the standard Nelder-Mead algorithm (Nelder and Mead 1965).

### SNP selection

The method assumes no LD. To accommodate this assumption, we use the following scheme: We divide the genome into 100 Kb regions and choose just one SNP for each region - the SNP with allele frequency closest to 0.5 in the reference population. We note that other schemes could be used to identify the most informative set of SNPs that are in approximate linkage equilibrium with each other. However, we found that more aggressive schemes explored, that quantify the information in the SNPs using the observed suspect or victim genotypes, tended to generate an elevated level of false positives.

### Testing for population structure

The method, in its simplest form, assumes that the reference panel is representative of the pool of individuals from which the suspect is sampled. If that is not the case, we would expect to observe false positives. Hence, we implemented as test to identify population subdivision between the reference panel and the suspect. For each SNP, let *f*_*i*_ be the major allele frequency. We then code the suspect genotype as its major-allele dosage *g*_*i*_ = 1.0 if homozygous major, 0.5 if heterozygous, 0.0 if no major allele and calculate the test statistic

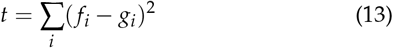

where the sum is over all valid SNPs. We generate a null distribution using simulations of 1,000 pseudo-suspects, by sampling genotypes from the panel frequencies assuming HWE and independence among SNPs. To quantify the overall degree of mismatch between the reference population and the suspect, we then calculate

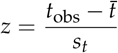

where 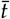 and *s*_*t*_ is the mean and standard deviation of *t* in the simulated data, and *t*_obs_ is the value of *t* in the observed data. A value of |*z* |> 2 then signifies a mismatch between suspect and reference population.

### Incorporating additional errors

So far we have assumed that errors can be modeled using genotype likelihoods. We note that the possibility of additional errors, not modeled by standard methods, can be incorporated in various ways by modifying read likelihood scores. A simple symmetric way of doing this is to modify the read nucleotide likelihood, *p*(*x*_*i*_|*nt* = *j*), *x*_*i*_ *∈ S*, to be

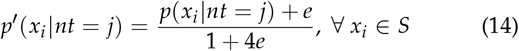

which is then included in the calculation of all terms of the likelihood function. This gives a possibility of evaluating the strength of the evidence if the error rate is different from that assumed in initial calculations.

### Simulations

To test the proposed method we turned to simulations. We simulated under the following scenarios: (1) Presence (or absence) of the suspect, victim and contamination from the general population in varying proportions. (2) Presence (or absence) of the suspect, victim and contamination from a single person sampled from the general population in varying proportions. (3) Presence of a family member of the suspect, the victim and contamination from the general population in varying proportions.

We simulated mixtures generated from phased 1000 Genomes Project genotypes. Unless otherwise stated, simulations used individual HG00100 as the true contributor, HG00110 as the known victim, and European (EUR) allele frequencies. Only biallelic single-nucleotide variants with a frequency between 0.01 and 0.99 in the specified allele-frequency field were eligible. Mixtures containing an unrelated individual used HG00260 as that individual. Population contamination was generated by sampling each contaminant allele independently from the reference and alternative alleles according to their site-specific population allele frequencies.

Simulations that include relatives were performed using the pedigree given in Figure 1. Notice the proband labeled “self” which typically will represents the contributor to the mixture in the simulations. Related genotypes were generated with ped-sim Caballero *et al*. (2019) from British (GBR) 1000 Genomes founder haplotypes 1000 Genomes Project Consortium *et al*. (2015). We used the sex-specific refined genetic map of Bhérer et al. Bhérer *et al*. (2017) and the crossover-interference parameter file distributed with ped-sim, which contains Housworth–Stahl interference-escape parameters estimated by Campbell et al. Campbell *et al*. (2015). The six-generation family was simulated for the true contributor, and the victim profiles were generated independently from that. Combined mixtures were then generated using the designated contributor, the unrelated victim, and population contamination in proportions 0.25, 0.50, and 0.25, respectively. The simulation used 10 *×* mean depth and approx. 27k SNPs (one in each 100kb genomic region), EUR frequencies, and a per-base error rate of 0.0002. L1 was calculated for 34 candidate profiles, including the true contributor, relatives ranging from first-degree relatives to second cousins twice removed, and six unrelated individuals.

**Figure 1.**
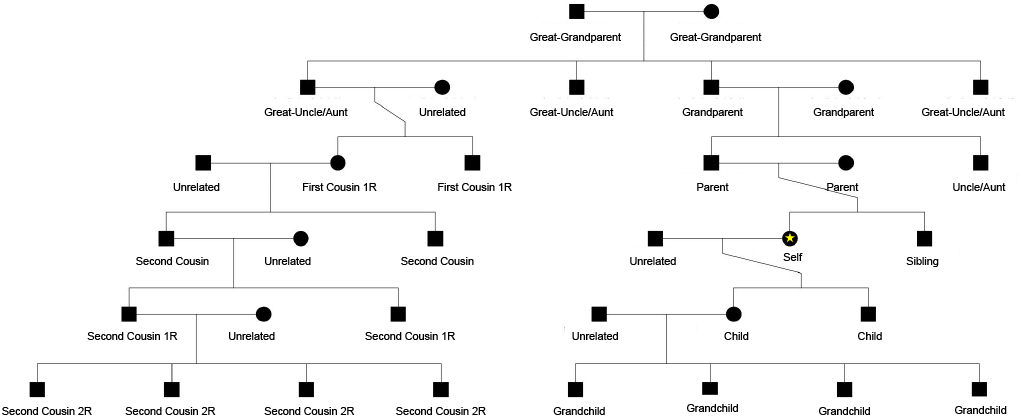
Simulated pedigree used to generate related candidate profiles. The true contributor to the mixture is labeled “Self”, and all relatives are labeled depending on their relation to that individual

At each locus, the number of reads contributed by source *j* was sampled independently from a Poisson distribution with mean *D f*_*j*_, where *D* is the requested mean sequencing depth and *f*_*j*_ is the source fraction. In heterozygous individuals, each allele was sampled with equal probability. The simulated reads from the different sources were merged into one file with a randomized order of the reads. Sequencing errors were generated by drawing a number of erroneous bases from a binomial distribution and replacing each selected base uniformly with one of the other three nucleotides with equal probability. Except for the sequencing-error experiment, the per-base error rate was 0.0002 and all observations were assigned Phred quality Q37.

The genome was divided into 100kb windows, and within each window, the locus with allele frequency closest to 0.5 was retained for mixture analyses, subject to the experiment-specific maximum number of SNPs. A separate set of one randomly chosen marker per window was used to test divergence between the suspect and the reference population. The crime-scene pileup, candidate and victim genotypes, and the corresponding allele frequencies were combined in JSON format for likelihood analysis. Workflows were managed with Snakemake; VCF files were sorted and indexed with bcftools, likelihood calculations were performed by the C implementation of DNAMIXTURE, and results were summarized in R using tidyverse.

## Results

We first evaluate Equation (5) for a range of different scenarios where the contaminant is either a single individual or a composite of many individuals from the population. We compared contamination generated from a single individual with contamination generated by independent population sampling. The true-contributor fractions were 0%, 1%, 5%, 10%, 20%, 30%, 40%, and 50%. For each value, the remaining fraction was divided equally between the victim and contaminant. In the individual-contamination condition the contaminant was HG00260; in the population-contamination condition contaminant alleles were sampled using EUR frequencies. Data were generated at a 10*×* mean depth, with approx. 27,000 SNPs and a per-base error rate of 0.0002. Ten replicates were generated for every mixture and source type.

Each simulated dataset was analyzed twice, once with the population-contamination likelihood and once with the single-individual-contamination likelihood, yielding a full 2 *×* 2 comparison of the generating process and fitted model.

As can be seen from Figure 2, when the contaminant is from the general population, and we use the likelihood ratio assuming contamination is from the general population, the likelihood ratio generally performs well. Similarly, the estimated proportions are quite accurate. We note that when the suspect is not in the mixture, two times the log likelihood ratio is reasonably well-fitted by a 50:50 mixture of a 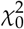 and 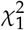 distribution as expected from asymptotic theory ((Self and Liang 1987) - see Supplementary Figure 1). Also, even at 5% mixture of the suspect, the contribution of the suspect can be detected. We observe similar performance when simulating and analyzing assuming the contamination is from a single individual. However, when using a mis-specified likelihood we find a high false-positive rate, particularly when using the individual analysis model and contamination is truly from the population. This suggests that the more conservative approach is to use the population contamination model, and we recommend that use, if using only a single model for analyses. We will discuss an approach this uses both models in a combined statistic below.

**Figure 2.**
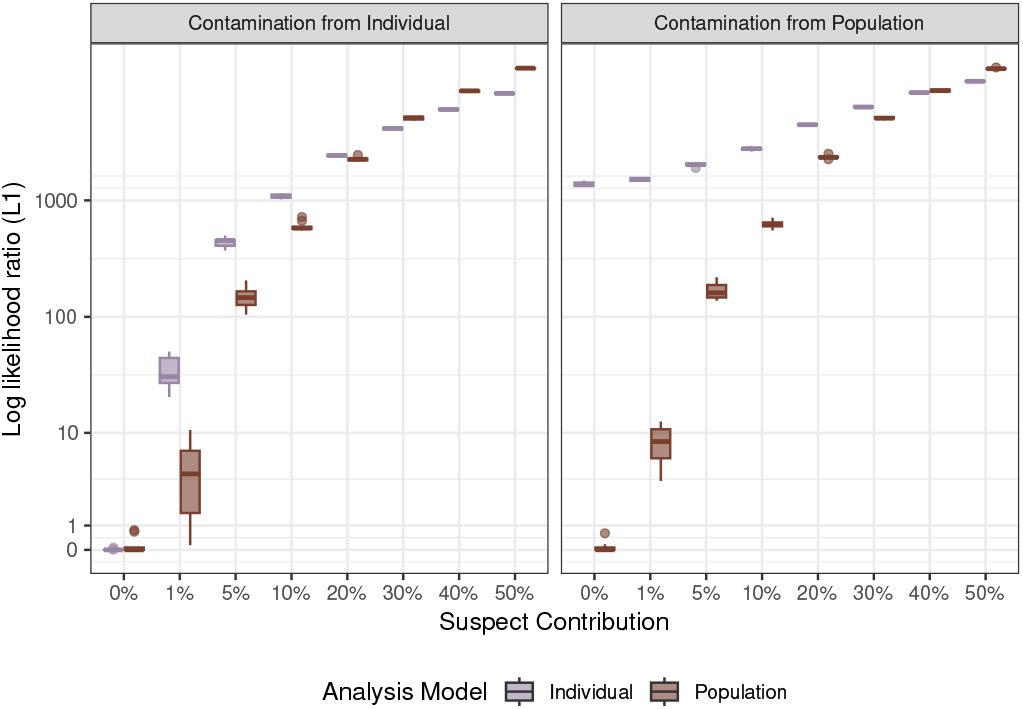
Log-likelihood ratios (*L*_1_) under alternative contamination scenarios. Boxplots show the distribution of log-likelihood ratios across simulation replicates for different levels of suspect DNA contribution. Results are shown separately for simulations assuming contamination from a single individual (left) and from a population (right). Within each panel, LRs are computed under two analysis models: an Individual model and a Population model.

We next (Figure 3A) evaluate the performance of Equation (5) when the suspect is absent from the mixture but a relative is present in the mixture. Notice, that large likelihood ratios are obtained even for relatives as distant as second cousins twice removed. This illustrates the risk of using likelihood ratios, or other statistical evidence, that does not evaluate the hypothesis that the true contributor to the mixture is a distant relative to the suspect. A large contribution from a relative might provide false evidence of a small contribution from the suspect.

**Figure 3.**
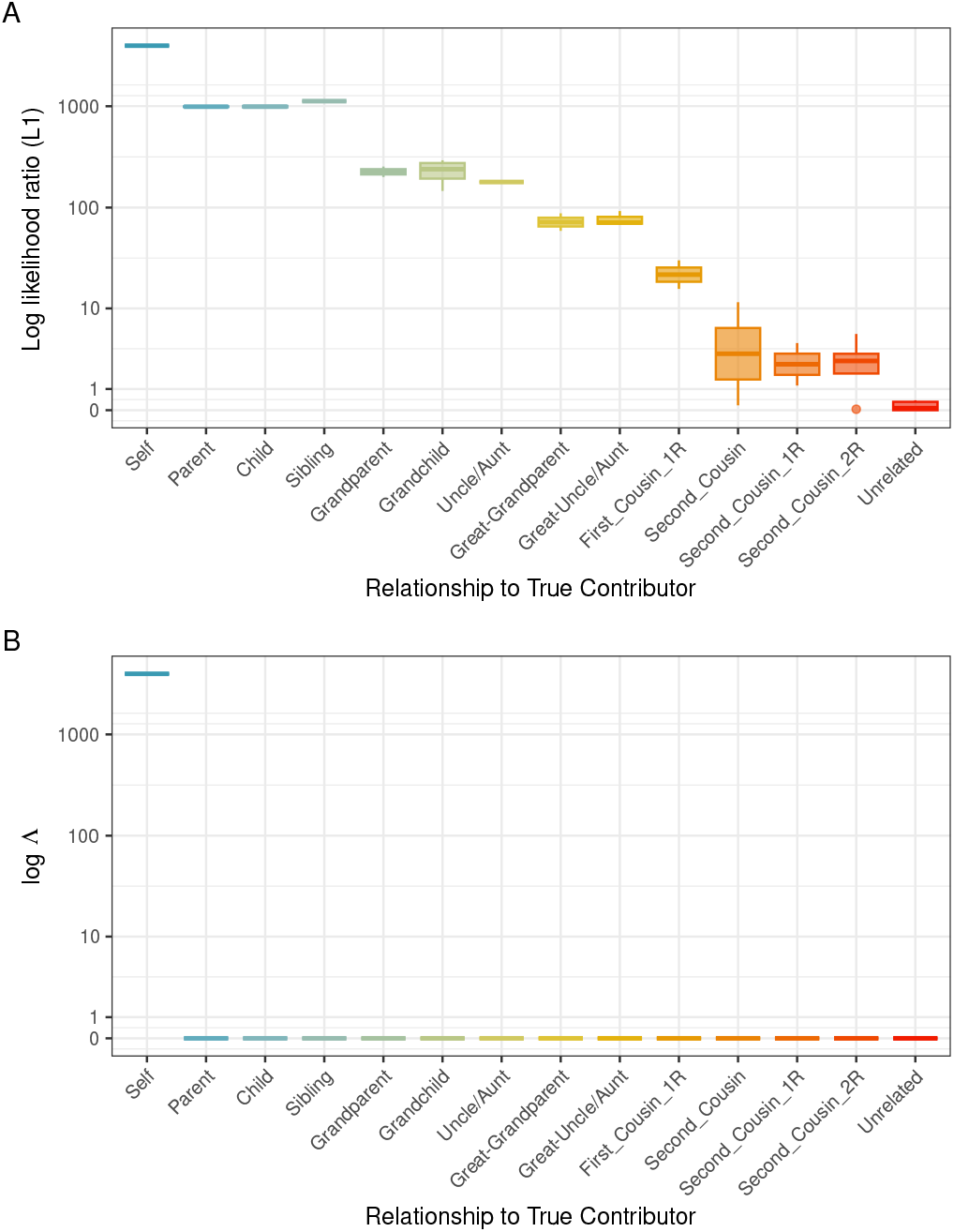
Log-likelihood ratios when the suspect (self) or a relative is present in the mixture. Boxplots show the distribution of *L*_1_ log-likelihood ratios among replicates for each genetic relationship to the true contributor. *L*_1_ values decrease with increasing genetic distance from the true contributor but remain elevated for close relatives, demonstrating the potential for false support when a relative of the suspect is the true contributor.

We then evaluated how well likelihood ratios comparing hypotheses positing a contribution from a relative versus a direct contribution from the suspect, are able to distinguish between these possible sources of the DNA (Figure 4, Supplementary Figure 2). Using the same pedigree mixtures as in Figure 3, we compared all related and unrelated candidates under *L*_1_, *L*_2_, *L*_3_, and *L*_4_. *L*_4_ was evaluated with (*k* = 1, 2, 3), corresponding to first-, second-, and third-cousin alternatives.

**Figure 4.**
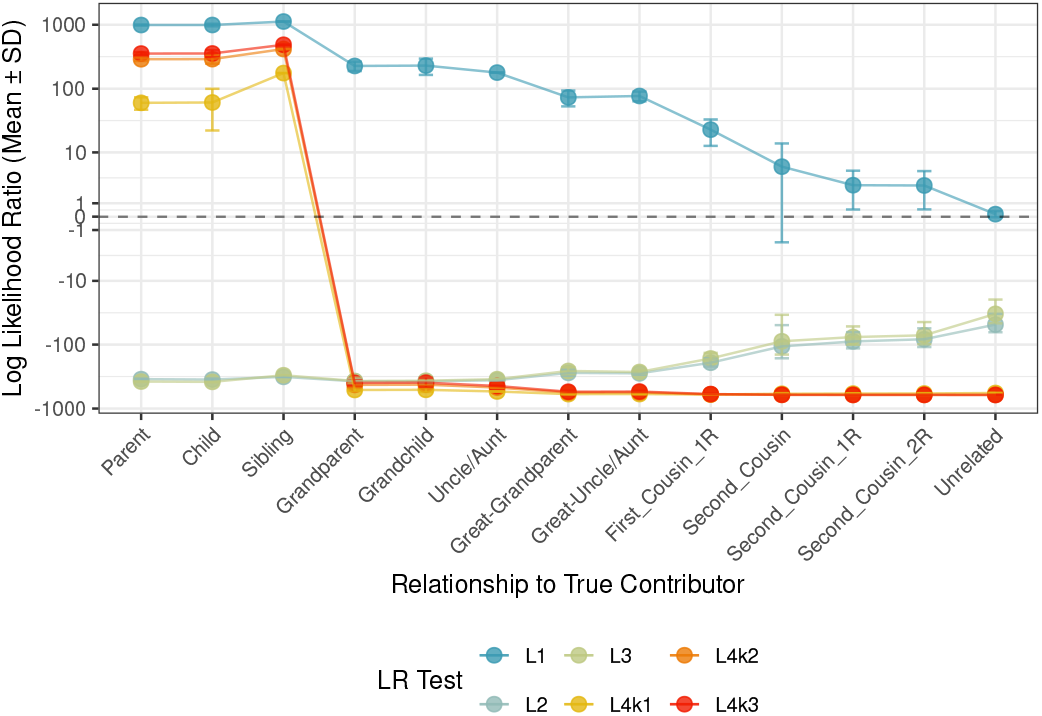
Comparison of likelihood ratio tests (*L*_1_, *L*_2_, *L*_3_ and *L*_4_ *k*_1_–*L*_4_ *k*_3_) for distinguishing the true contributor from related candidates. Each point shows the mean log-likelihood ratio across replicates for a given relationship to the true contributor; error bars show one standard deviation. Simulation conditions are as in Figure 3. *L*_1_ remains elevated for close relatives, whereas *L*_2_ (suspect vs. hypothesized sibling) and *L*_3_ (suspect vs. hypothesized parent) reduce or eliminate false support for first-degree relatives..

For each relationship class other than the true contributor, we calculated the mean and standard deviation of the log likelihood ratio for each test. Notice that, as in Figure 3A, the *L*_1_ statistic is misleading and reports large likelihood ratios when relatives, and not the suspect, are in the mixture. However, when likelihood ratios using the hypothesis of related individuals are used, they effectively distinguish between a contribution from the suspect and the relative. For example, using *L*_2_ or *L*_3_, the log likelihood ratio is on the order of −100 to −1000 when the true contributor is a parent, child, sibling, grandchild, grandparent, Uncle/Aunt, Great-Grandparent, or Great-Uncle/Aunt, when the likelihood ratio is defined such that small likelihood ratios favor a relative over the suspect.

These results in Figure 4 suggest a procedure that uses *L*_2_ or *L*_3_ in conjunction with *L*_1_. In particular, we propose reporting

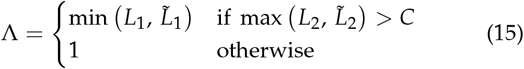

where *C* is a threshold that could be varied, but is set to *C* = 10 in the following analyses. Here, *L*_1_ and *L*_2_ are the likelihood ratios of Equations (5) and (7), respectively, computed under the general-contamination likelihood (Equation 2), and 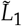 and 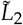 are the same ratios computed under the single-individual-contamination likelihood. That is, the smaller of the two suspect likelihood ratios is reported only if the evidence favors the suspect over a hypothesized sibling by at least a factor of *C* under at least one of the two contamination models; otherwise no evidence (Λ = 1) is reported. The hope is that this statistic will be robust to both the model mis-specification discussed for Figure 2, and protect against false identification of the suspect when a distant relative is in the mixture. As we will later show, this statistic also increases robustness to the presence of population structure.

The result of this procedure for protecting against model mis-specification is shown in Figure 5. Notice that there, by design, no longer are any false positives when using the individual contamination model to analyze data with contamination from the population. So an added benefit of this procedure is that it also protects against this type of false positives. The statistic offers a similar protection against false positives due to the presence of relatives (3B). When using this Λ, the false positives observed in 3A vanish.

**Figure 5.**
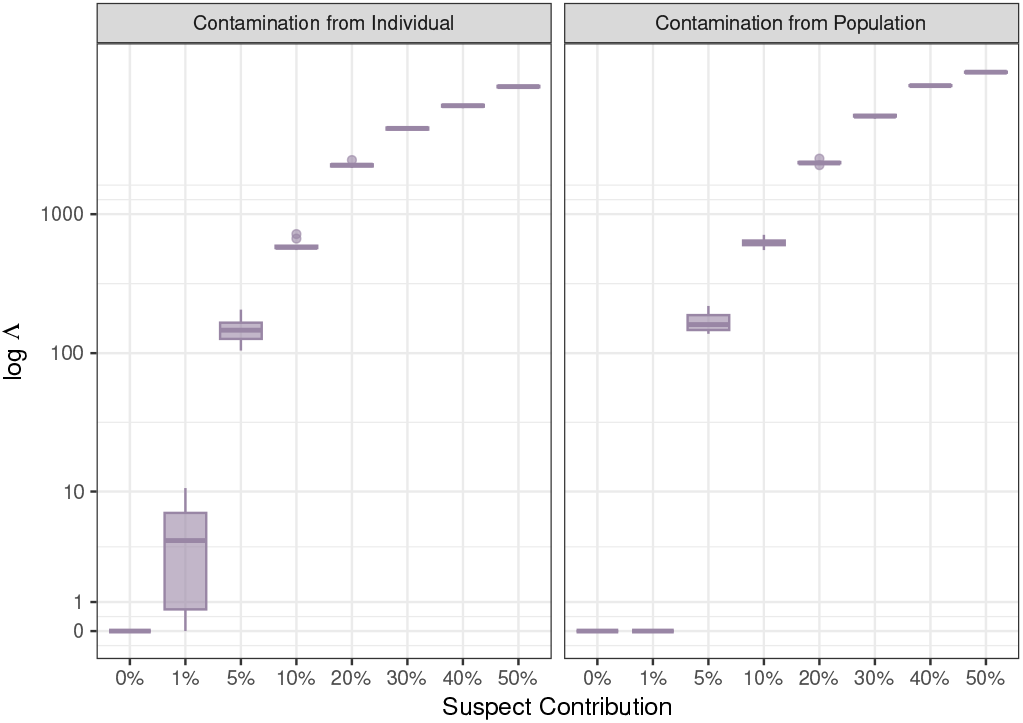
The combined evidence score, Λ, under alternative contamination scenarios. Calculations are done for Λ but the conditions are otherwise the same as in Figure 2.

We next examined the robustness of the statistics to population structure by using allele frequencies from ancestry panels that may not match the suspect well. A mixture containing 25% NA18561 (CHB; East Asian), 50% HG00110 (GBR; European), and 25% population contamination was simulated at 10*×* mean depth, with approx. 27,000 SNPs and a per-base error rate of 0.0002. Forty-two candidate individuals were sampled from 14 populations, with three individuals per population: ASW, CEU, CHB, CHS, CLM, FIN, GBR, IBS, JPT, LWK, MXL, PUR, TSI, and YRI. These represented four superpopulations (AFR, AMR, EAS, and EUR) and included the true contributor. Every candidate was analyzed with *L*_1_ (Figure 6A) and Λ (Figure 6B) using five alternative frequency definitions: global allele frequency and EAS, AMR, AFR, and EUR superpopulation frequencies. For each combination of suspect and reference population, we also performed the previously described test of population structure (ancestry-compatibility test; see also Supplementary Figure 3). Results were stratified by candidate superpopulation, true-contributor status, and the ancestry-compatibility test. Notice that substantial amount of false positives can be obtained when there is a mismatch between the suspect and the ancestry panel (Figure 6A). Strong inflation of likelihood ratios is observed for ancestry-mismatched individuals, particularly when allele-frequency panels poorly represent the suspect ancestry. Perhaps surprisingly, this is true, even when filtering based on the test of allele frequency mismatch between the suspect and the reference panel (Figure 6A; lower panel). In the case of an American suspect tested, when global allele frequencies are used, the test of population structure does not reject in several cases that yield high likelihood ratios in the *L*_1_ test. The individuals do not have allele frequencies that deviate from the global population more than expected from a randomly sampled individual, but the individual is still genetically closer to the true perpetrator than a randomly sampled individual and the *L*_1_ test is therefore significant. This is an example of the general problem of suspects and true perpetrators that are genetically closer to each other than two random individuals sampled from the population. However, and perhaps surprisingly, the use of Λ seems to fully protect against population structure in this setting, whether the test for population structure is carried out or not.

**Figure 6.**
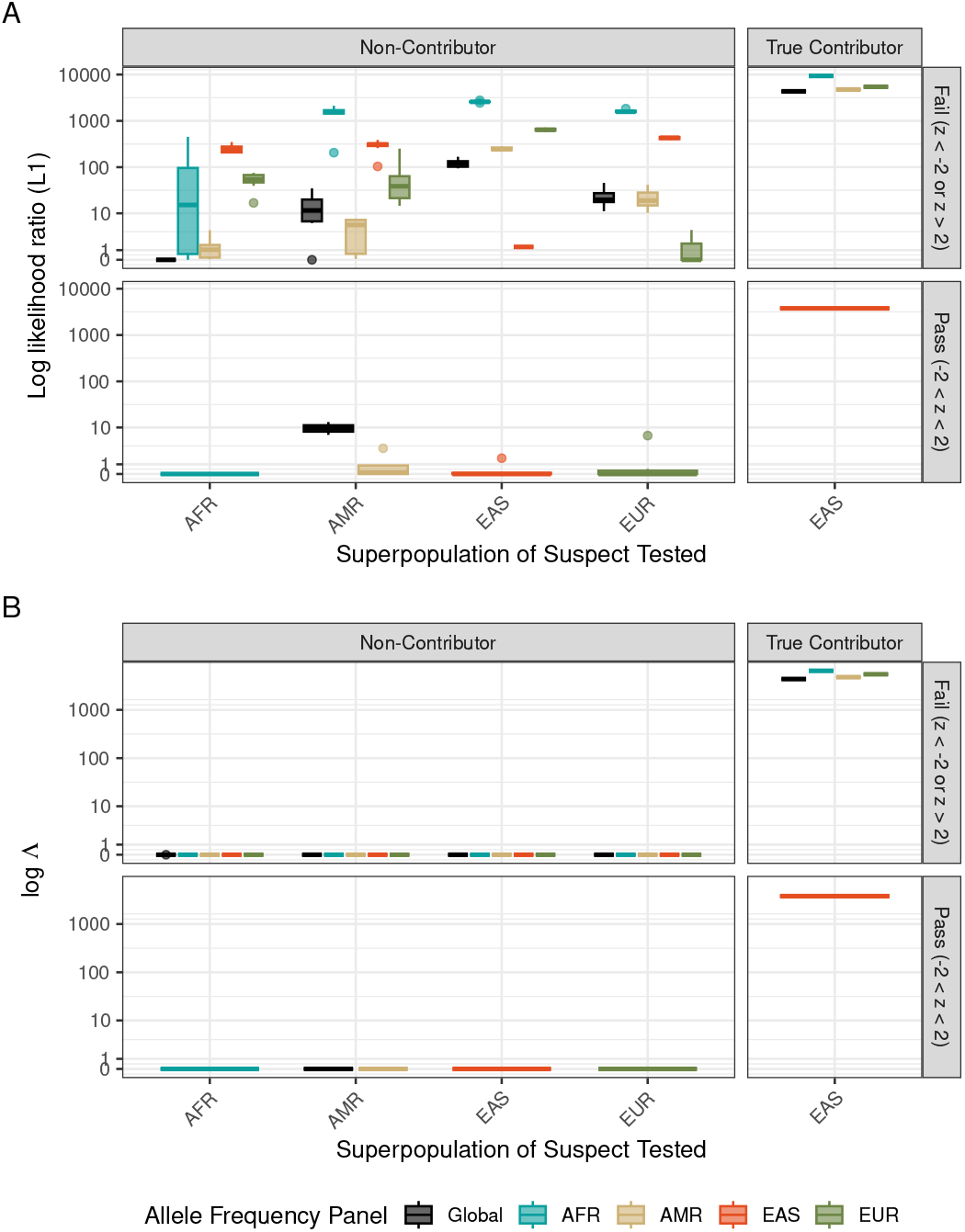
Robustness of likelihood ratios to ancestry mismatch between suspects and allele-frequency reference panels. Suspects were stratified according to genetic similarity to the reference population using the population stratification test (z-score threshold above 2 or below −2). Boxplots show log likelihood ratios obtained using different allele-frequency panels across suspect superpopulations. Panel A shows the results using the likelihood ratio test based on *L*_1_ and population contamination and Panel B shows the results using Λ

**Figure 7.**
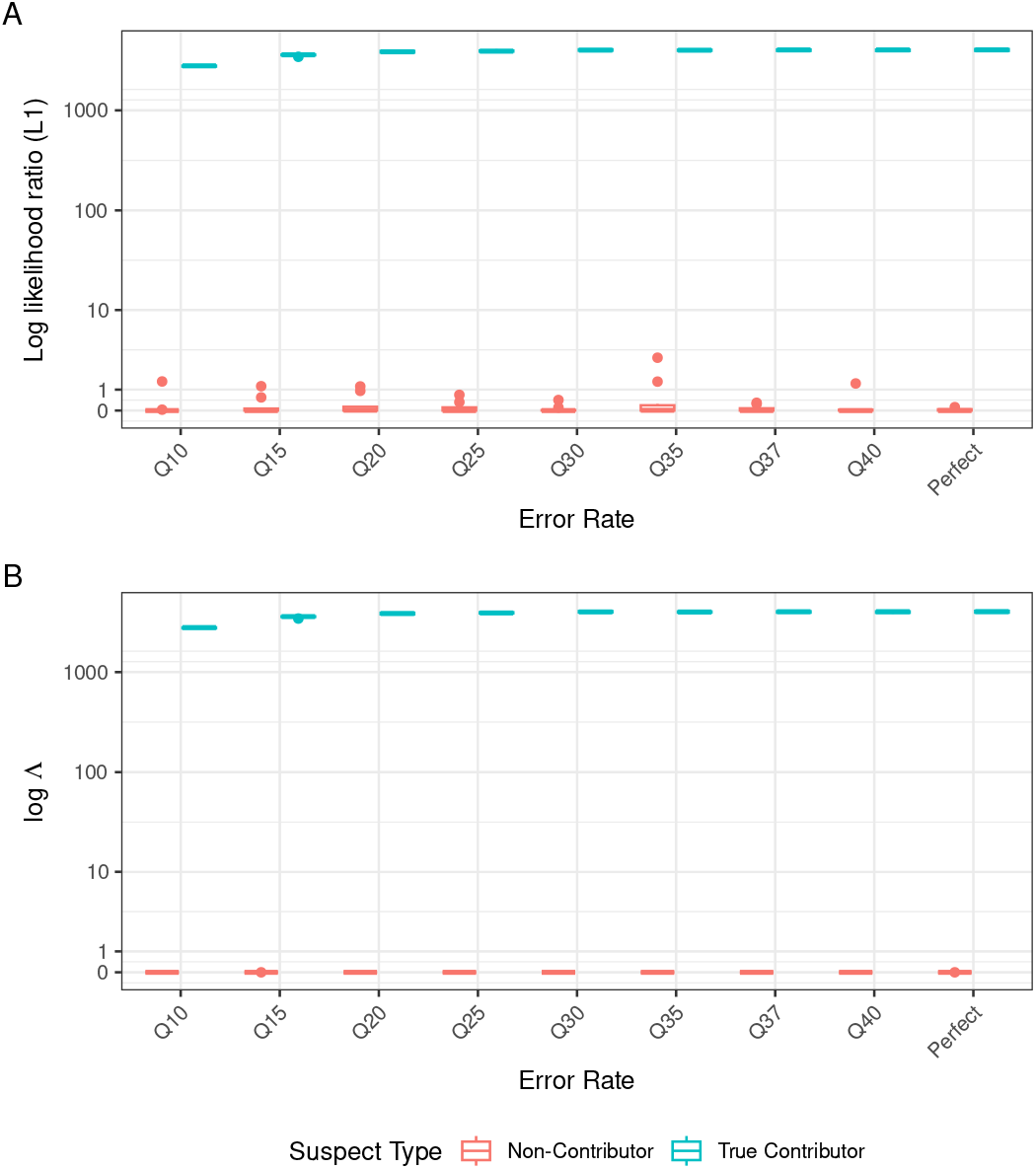
Robustness of likelihood ratios to sequencing error. The Figure shows the results for simulated mixtures comprising 25% true-contributor DNA, 50% victim DNA, and 25% population contamination at 10*×* mean depth and approx.

To test sensitivity to base-call error, we simulated data with varying error rates. All simulations assumed a reported quality of Q37 but the true error rate was varied; the experiment therefore measures robustness to discordance between the true error process and the quality score assumed by the likelihood calculation. Ten replicates were generated per error rate. HG00100 and HG00260 were tested using *L*_1_ with population contamination (Panel A) and using Λ (Panel B). As sequencing error increases (lower quality score), there is no evidence of an increase of false positives, but there is a slight reduction in power to the detect the true contributor when error rates are high. These results suggest that the method retains discriminatory power even under highly elevated sequencing error rates while maintaining low false-positive support for unrelated individuals not in the mixture. Among the many challenges the use of SNPs for DNA mixtures face, sequencing errors do not appear to be the most critical one.

We next evaluated robustness to sequencing depth (Figure 8). We jointly varied mean sequencing depth, number of SNPs, and mixture composition (Figure 8); Supplementary Figures 4 and 5). Notice that low number of SNPs or low sequencing depth does not cause an excess of false positives when using Λ. However, there is naturally reduced power to detect the suspect when there is low sequencing coverage or few SNPs. Nonetheless, a suspect contribution of 50% can reliably be identified at 1 *×* depth with 1,000 SNPs.

**Figure 8.**
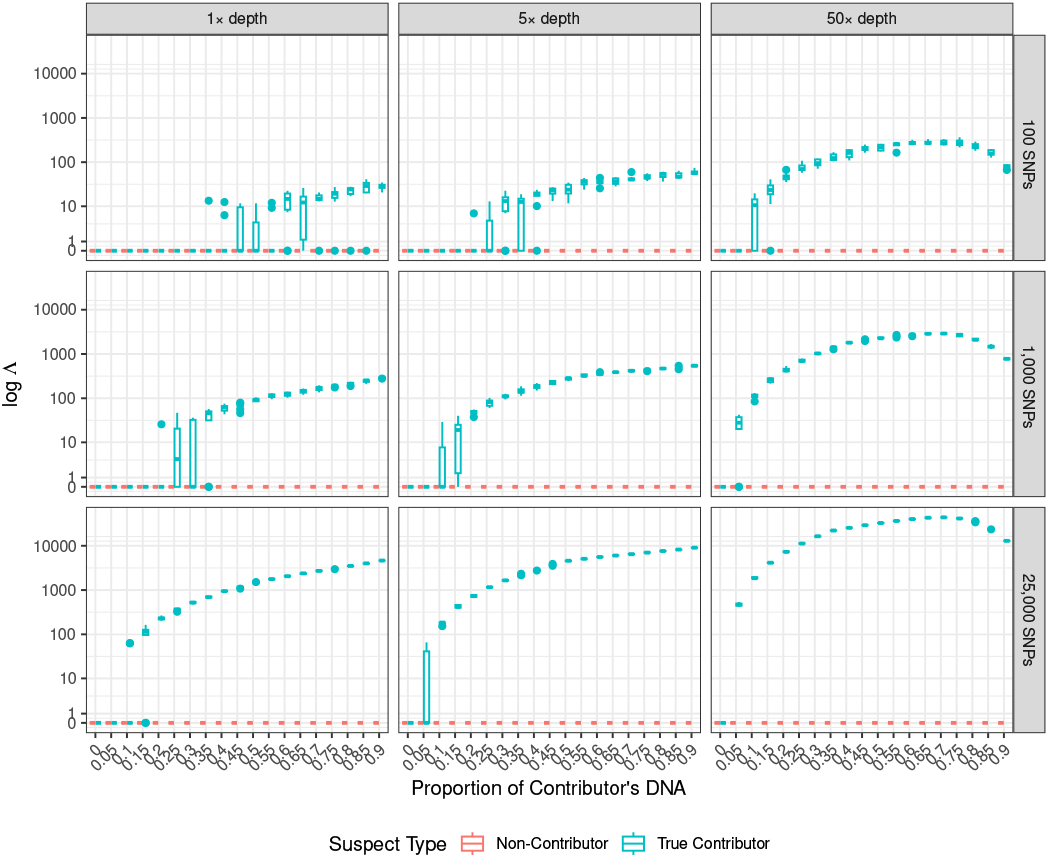
Λ as a function of sequencing depth and marker number. Boxplots show the distribution of log Λ across ten simulation replicates as a function of the true contributor’s proportion of the mixture. Mixtures contained DNA from the true contributor (HG00100) at the indicated proportion, victim DNA (HG00110) at a fixed proportion of 10%, and population contamination simulated from EUR allele frequencies making up the remainder, with a per-base error rate of 0.0002. In each replicate, both the true contributor and a non-contributor (HG00260) were tested as the suspect. Panels show combinations of mean sequencing depth (columns: 1*×*, 5*×*, 50 *×*) and number of markers (rows: 100, 1,000, 25,000). Results for the full grid of depths (1–100 *×*) and marker numbers (100–25,000), and for the corresponding raw *L*_1_ log likelihood ratios, are shown in Supplementary Figures 4 and 5.

We assessed the effect of physical marker spacing with and without prior linkage-disequilibrium (LD) pruning. Variants were pruned with PLINK `--indep-pairwise 50 5 0.2`. Mixtures contained 25% HG00100, 50% HG00110, and 25% population contamination, with 10× mean depth and a per-base error rate of 0.0002. From both the unpruned and LD-pruned inputs, up to 2,000 mixture markers were selected using 27,000 SNPs. Boxplots show the distribution of log likelihood ratios across simulation replicates for a true contributor (blue) and a non-contributor (red) under increasing sequencing error rates corresponding approximately to Phred quality scores ranging from perfect (no error) to Q10. Per-base error probabilities were 0.10000, 0.03162, 0.01000, 0.00316, 0.00100, 0.000316, 0.000200, 0.000100, and 0, corresponding to Q10, Q15, Q20, Q25, Q30, Q35, Q37, Q40, and error-free data, respectively. Panel A shows the results using the likelihood ratio test based on *L*_1_ and population contamination and Panel B shows the results using Λ non-overlapping physical windows of 1, 5, 10, 50, 100, 500, 1,000, 5,000, 10,000, 50,000, 100,000, 500,000 bp. Within each window the marker with allele frequency closest to 0.5 was retained. Ten replicates were generated per condition. HG00100 and HG00260 were analyzed with HG00260 representing a potential false positive.

Perhaps surprisingly, there is not much effect of LD pruning (Figure 9). In the presence of LD, SNPs are correlated and we can consider the likelihood function to be a composite likelihood. As such, we would expect the log likelihood ratio to have the same expectation as when SNPs are independent, but possibly a different variance (see e.g., Varin *et al*. (2011)). It is, therefore, maybe not too surprising that likelihood ratio still is able to distinguish between the suspect and other individuals and does not, at least in these limited simulations, tend to produce false positives. However, the probabilistic interpretation of the likelihood ratio changes. Also, we cannot exclude the possibility that in many other scenarios, there would be a larger effect of LD pruning. We, therefore, recommend the use of approximately independent SNPs, for example identified by simply choosing just one SNP, with high minor allele frequency, in each 100kb region of the genome, as in this study.

**Figure 9.**
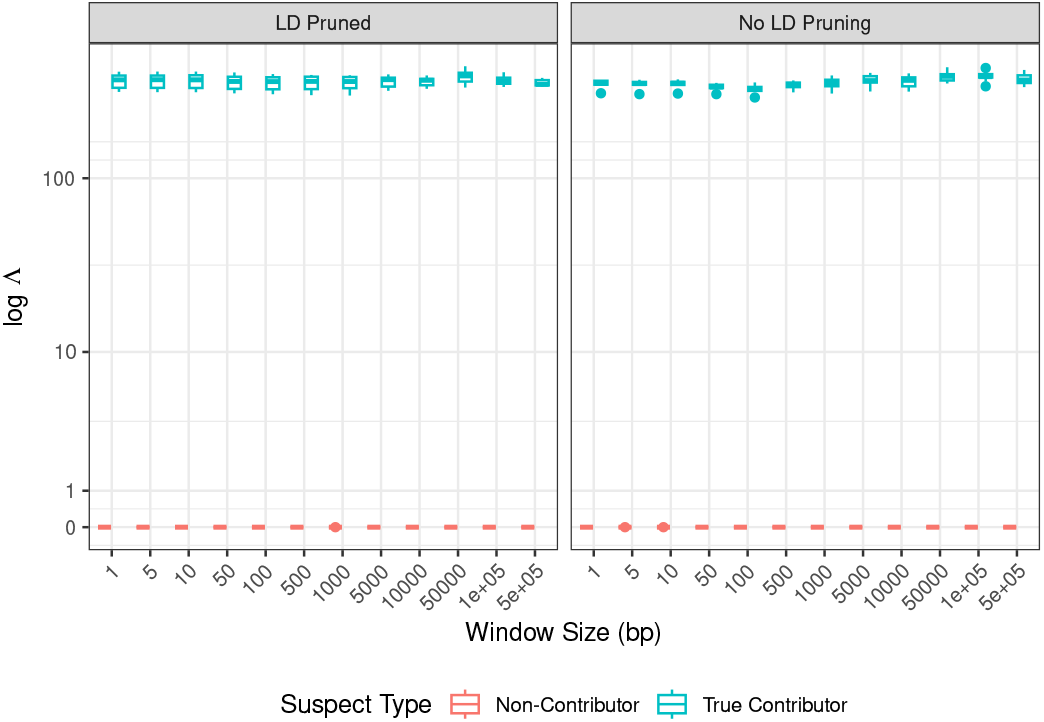
Effect of linkage disequilibrium pruning on likelihood ratio performance across SNP selection window sizes. Log likelihood ratios (LRs) are shown for a true contributor (HG00100, blue) and a non-contributor (HG00260, red) across increasing SNP selection window sizes (1–500,000 bp), with and without LD pruning applied to the reference panel. Simulated mixtures consisted of 25% contributor, 50% victim, and 25% contamination, sequenced at 10× coverage. For each window size, 2,000 SNPs were selected and LRs computed across replicates.

## Discussion

The use of shotgun sequencing for source identification of mixed DNA stains has great promise as a tool for forensic analyses. However, the interpretation of the data produced by such sequencing is challenged by sequencing errors, population structure, the potential presence of related individuals, linkage and linkage disequilibrium. In this paper, we have derived and implemented statistical methods for identifying the sources in a mixture, but we have also shown that likelihood ratios can be unreliable if the wrong model of contamination is used and/or if there is unmodeled population structure or related individuals in the mixture. We have not attempted in this paper to address the problem of population structure by modeling excess identity by descent between the suspect and an otherwise unrelated perpetrator. Classical methods use an *F* factor for forensic calculations of match probabilities (Balding and Nichols 1994) for this. However, in preliminary work we did not succeed in constructing a robust method that worked well under the assumption of both individual and population contamination, although we do not exclude the possibility that such a method could be constructed. Even so, such a method would still struggle with the problem of related individuals. Instead, we propose to increase robustness by using the statistic Λ which is the minimum of the log likelihood ratios calculated under the individual and population contamination models when at least one of the likelihood ratios favor presence of the suspect over the suspect’s sibling with a likelihood ratio > 10, and otherwise 0 (Equation 15). The use of this procedure appears to protect against several factors that otherwise could lead to false positives, such as population structure, and our recommendation is that this statistic should be the one presented to summarize the evidence that the suspect is in the mixture. We also recommend additional use of the test for genetic differentiation between the suspect and the reference panel. However, we note that Λ has only been tested in the few scenarios explored in this paper. There could be other conditions, such as other complex mixtures of contamination, under which the performance of Λ degrades. Also, SNP selection may affect the results. We here chose a SNP selection procedure in which the allele with frequency closest to 50% was chosen in non-overlapping genomic windows. We note that SNP data selected using other procedures might introduce new biases that are not well-modeled by our set-up. For these reasons, we do not recommend using Λ as stand-alone evidence in court.

## Data availability

The likelihood ratios and the Λ evidence score is implemented in the C program DNAMIXTURE, the C program available at https://github.com/ras-nielsen/DNAMIXTURE under the CC BY-NC-SA 4.0 license. Documentation, including the input format and a worked example, is at https://ras-nielsen.github.io/DNAMIXTURE/.

## Acknowledgements

We thank Lasse Vinner for discussions. This research was supported by NIH grant R35GM153400.

