## Supplemental Figures for "Forensic likelihood ratios for shotgun sequencing data from DNA mixtures"

September 28, 2026

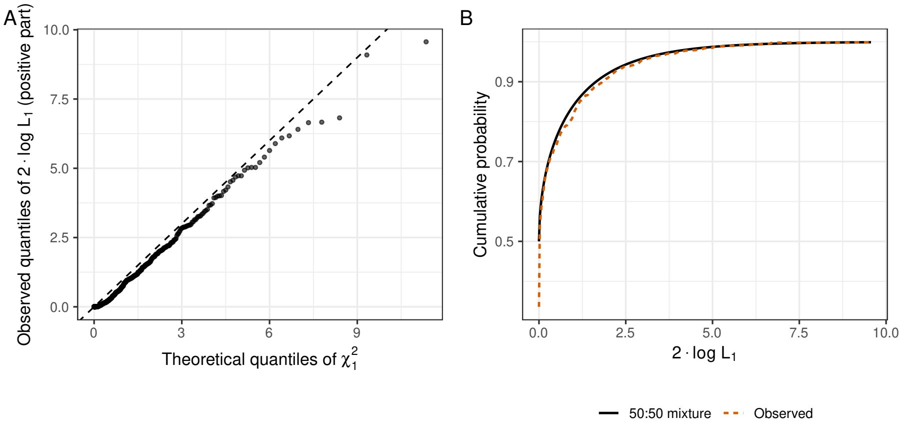


Figure 1: Null calibration of the $L_1$ test statistic. Two times the log likelihood ratio, $2\log L_1$, was computed for 1,000 simulated mixtures containing no DNA from the tested suspect (HG00260). Each mixture consisted of 50% victim DNA (HG00110) and 50% contamination sampled from EUR population allele frequencies, simulated at 10× mean depth with approximately 27,000 SNPs and a per-base error rate of 0.0002, and analyzed under the population contamination model, matching the simulation. Because the suspect contribution is estimated on its boundary ($f_1 = 0$) under the null hypothesis, the asymptotic distribution of $2\log L_1$ is an equal mixture of a point mass at zero and a $\chi^2(1)$ distribution (Self and Liang 1987). (A) Quantiles of the positive values of 2 $\log L_1$ plotted against $\chi^2(1)$ quantiles; the dashed line is the identity. (B) Empirical cumulative distribution of $2\log L_1$ (black) compared with the theoretical 50:50 mixture (orange). 52% of replicates fall at zero, consistent with the expected 50%.

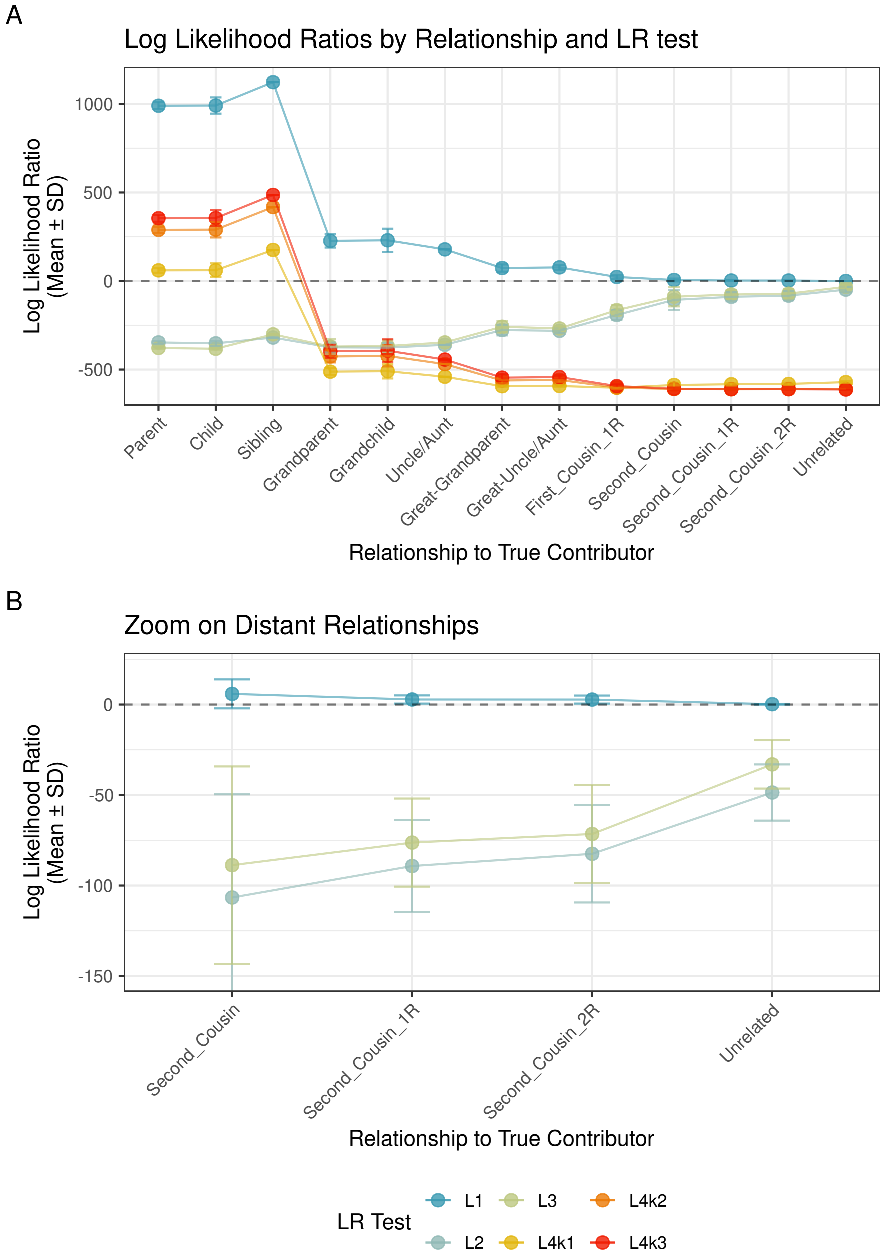
A
Log Likelihood Ratios by Relationship and LR test
Log Likelihood Ratio (Mean ± SD)
1000
500
0
-500
Parent
Child
Sibling
Grandparent
Grandchild
Uncle/Aunt
Great-Grandparent
Great-Uncle/Aunt
First_Cousin_1R
Second_Cousin
Second_Cousin_1R
Second_Cousin_2R
Unrelated
Relationship to True Contributor
B
Zoom on Distant Relationships
Log Likelihood Ratio (Mean ± SD)
0
-50
-100
-150
Second_Cousin
Second_Cousin_1R
Second_Cousin_2R
Unrelated
Relationship to True Contributor
LR Test
L1
L2
L3
L4k1
L4k2
L4k3

Figure 2: Log likelihood ratios by relationship to the true contributor on a linear scale. Simulations and likelihood ratio tests are identical to Figure 4, only the presentation differs, with the y-axis shown on a linear rather than a logarithmic scale. (A) All relationships. (B) Zoom on second cousins (including once and twice removed) and unrelated individuals. The $L_4$ statistics fall below the plotted range in this panel.

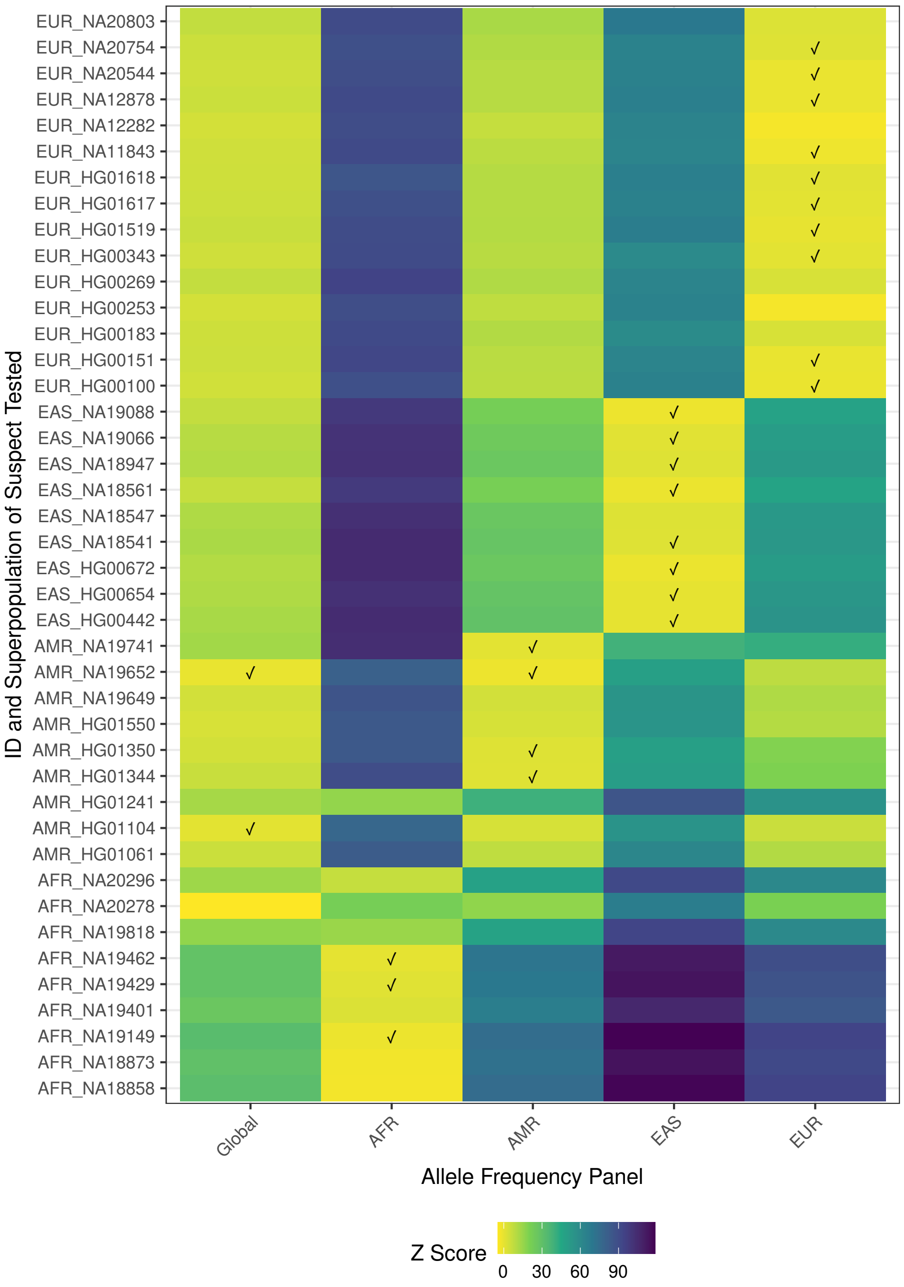

ID and Superpopulation of Suspect Tested
EUR_NA20803
EUR_NA20754
EUR_NA20544
EUR_NA12878
EUR_NA12282
EUR_NA11843
EUR_HG01618
EUR_HG01617
EUR_HG01519
EUR_HG00343
EUR_HG00269
EUR_HG00253
EUR_HG00183
EUR_HG00151
EUR_HG00100
EAS_NA19088
EAS_NA19066
EAS_NA18947
EAS_NA18561
EAS_NA18547
EAS_NA18541
EAS_HG00672
EAS_HG00654
EAS_HG00442
AMR_NA19741
AMR_NA19652
AMR_NA19649
AMR_HG01550
AMR_HG01350
AMR_HG01344
AMR_HG01241
AMR_HG01104
AMR_HG01061
AFR_NA20296
AFR_NA20278
AFR_NA19818
AFR_NA19462
AFR_NA19429
AFR_NA19401
AFR_NA19149
AFR_NA18873
AFR_NA18858
Global
AFR
AMR
EAS
EUR
Allele Frequency Panel
Z Score
0
30
60
90

Figure 3: Population-match z scores by suspect and allele frequency panel. Mixtures contained 25% DNA from the true contributor (NA18561, EAS), 50% from the victim (HG00110), and 25% population contamination simulated from EUR allele frequencies, at 10× mean depth with approximately 27,000 SNPs and a per-base error rate of 0.0002. Each of 42 suspects sampled from the AFR, AMR, EAS, and EUR superpopulations of the 1000 Genomes Project (rows) was analyzed using five allele frequency panels (columns: global, AFR, AMR, EAS, EUR). For each analysis, the population-match statistic, $t$, was computed from the suspect's genotypes at an independent set of approximately 27,000 background SNPs (one randomly chosen site per 100 kb window). The resulting $z$ score compares the observed value of $t$ to its distribution across 1,000 genotype vectors simulated indendepently under Hardy–Weinberg equilibrium from the panel frequencies. Check marks indicate $| z |< 2$, i.e., panels accepted as consistent with the suspect's ancestry.

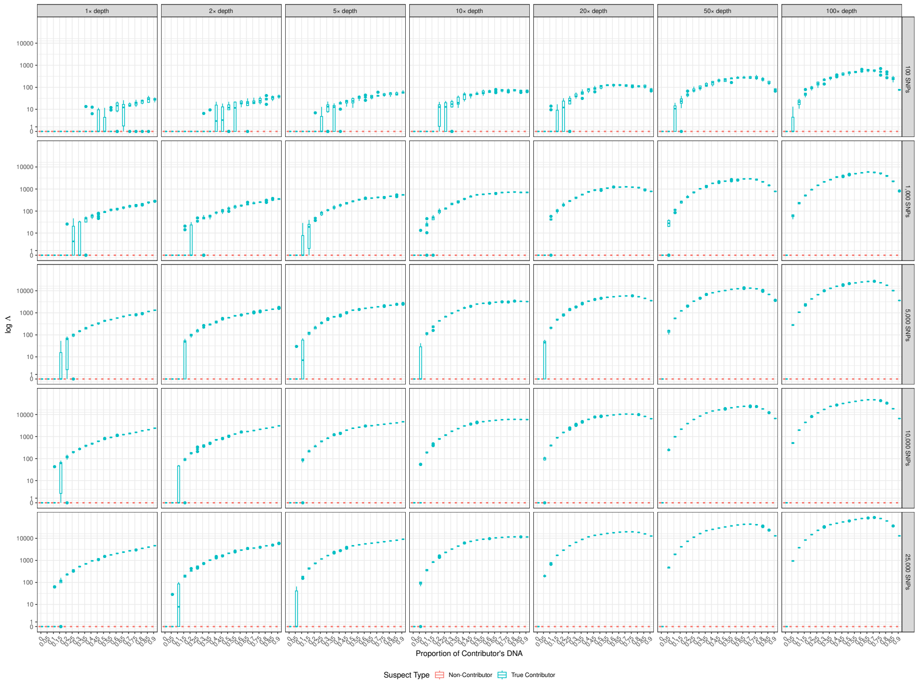


Figure 4: Λ for all combinations of sequencing depth and marker number. Simulations are identical to Figure 8, which shows a subset of these panels; here all combinations of mean sequencing depth (columns: 1×, 2×, 5×, 10×, 20×, 50×, 100×) and number of markers (rows: 100, 1,000, 5,000, 10,000, 25,000) are shown.

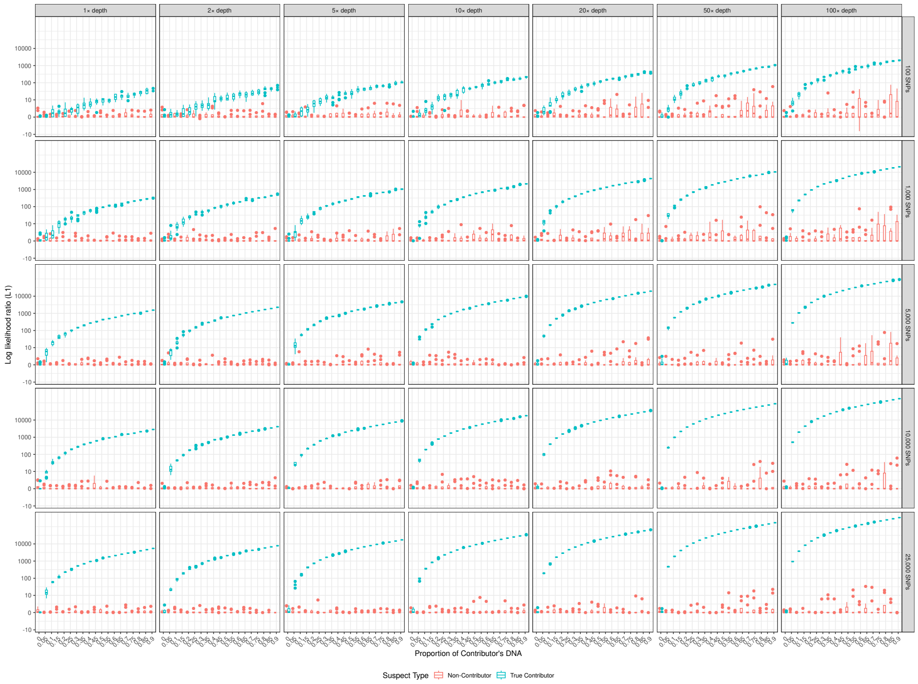


Figure 5: $L_1$ log likelihood ratios for all combinations of sequencing depth and marker number. Identical to Supplementary Figure 4, except that the y-axis shows the $L_1$ log likelihood ratio computed under the population contamination model.
